# A WAX INDUCER1/SHINE transcription factor controls cuticular wax in barley

**DOI:** 10.1101/2022.03.11.483819

**Authors:** Trisha McAllister, Chiara Campoli, Mhmoud Eskan, Linsan Liu, Sarah M. McKim

**Affiliations:** Division of Plant Sciences, University of Dundee

**Keywords:** barley, cuticle, *eceriferum*, wax, SHINE transcription factor

## Abstract

All land plants seal their above ground body parts with a lipid-rich hydrophobic barrier called the cuticle that protects tissues from dehydration and other terrestrial threats. Mutational studies in several model species, including barley, have resolved multiple loci regulating cuticular metabolism and development. Of particular importance are the *eceriferum* (*cer*) mutants characterized by visual alterations in cuticular wax. In barley, some *cer* mutants, such as *cer-x* lines, show defects in the distinctive β-diketone-enriched wax bloom on reproductive stage leaf sheaths, stems and spikes. In our study we exploited extensive allelic populations, near-isogenic lines and powerful genotyping platforms to identify variation in the *HvWAX INDUCER1* (*HvWIN1*) gene as causal for *cer-x* mutants. We show that HvWIN1 function reduces cuticular permeability, promotes the accumulation of β-diketones, and regulates cuticular metabolic gene expression. Analyses across the barley pangenome and hundreds of exome-capture datasets revealed high sequence conservation of *HvWIN1* but also two non-synonymous variants exclusive to cultivated germplasm. Taken together, we suggest that variation in *HvWIN1* controls multiple cuticular features in barley by controlling the expression of genes involved in cuticle development.

## Introduction

Land plants have a waxy, reflective cuticle to protects them from dangers of terrestrial life, including desiccation, pathogen attack and UV damage (Jeffree, 2006; Yeats and Rose, 2013). The cuticle forms on epidermal plant tissues exposed to the atmosphere and consists of a matrix, made up of the lipid biopolymer cutin embedded with polysaccharides and intracuticular waxes, which is covered by an epicuticular wax layer (Shepherd and Wynne Griffiths, 2006; Yeats and Rose, 2013; Guzmán *et al.*, 2014). Epicuticular waxes are mostly composed of acyl aliphatics arranged as films or crystals which can impart a glossy or glaucous appearance, respectively (Yeats and Rose, 2013). Biochemical and genetic studies showed that cuticular waxes originate from the endoplasmic reticulum where Fatty Acid Elongase complexes, including β-ketoacyl-CoA synthases (KCSs), elongate C_16_ or C_18_ fatty acids (FAs) into very-long-chain fatty acids (VLCFAs, > 20 carbons) which are either recruited into the decarbonylation pathway, making odd-numbered alkanes, secondary alcohols and ketones, or into the alcohol forming pathway, producing even-numbered primary alcohols and wax esters (Kolattukudy PE, 1976; Samuels *et al.*, 2008; Yeats and Rose, 2013). While most land plant cuticles share these compounds, exact cuticle composition and structure vary widely depending on tissue, species, developmental stage and environmental conditions, which likely reflect adaptive responses to diverse niches and growth habits (Jetter *et al.*, 2006; Buschhaus and Jetter, 2011; Edwards and Kenrick, 2015; Gosney *et al.*, 2016; Xue *et al.*, 2017; Domínguez *et al.*, 2017). For instance, grasses, including our cereal staples, show cuticular properties which may help these plants to cope with drought-prone environments, and accordingly, influence grain yields (Dodd and Poveda, 2003; Zhang *et al.*, 2013*a*; Guo *et al.*, 2016; Bi *et al.*, 2017; Xue *et al.*, 2017; Laskoś *et al.*, 2021). In particular, the distinctive blue-green glaucous wax bloom on exposed leaf sheaths, stem nodes and internodes, and inflorescences during the reproductive phase of many graminoid crops may provide protection from pests, UV damage and water loss (Tulloch and Hoffman, 1980; Richards *et al.*, 1986; Febrero *et al.*, 1998; González and Ayerbe, 2010; Zhang *et al.*, 2013*a*; Laskoś *et al.*, 2021) and may have been selected accordingly during barley and wheat domestication (Hen-Avivi *et al.*, 2016; Nice *et al.*, 2016).

To learn more about the genetic and developmental basis of the wax bloom, we study cuticular mutants in barley. Many of these are *glossy* or *eceriferum (cer)* mutants, identified in mutagenesis screens for reduced glaucousness (Lundqvist and Wettstein, 1962). To date, researchers have identified the genes underlying several *cer* mutants, including *Hv3-KETOACYL-COA SYNTHASE 1* (*HvKCS1*) and *Hv3-KETOACYL-COA SYNTHASE 6* (*HvKCS6*) (Weidenbach *et al.*, 2014; Li *et al.*, 2018). In addition, cloning of the *CER-CQU* metabolic gene cluster revealed the genes necessary for production of the C_31_ β-diketone crystalline tubes comprising the main component of the barley and wheat epicuticular wax bloom. The cluster encodes three enzymes – a diketone synthase (DKS), a lipase/carboxyl transferase, and a P450 hydroxylase responsible for adding hydroxyl (OH) groups onto β-diketones – which underly the barley *Cer-C, Cer-Q* and *Cer-U* loci respectively, and whose orthologs are also present within the *W1* metabolic gene cluster mediating β-diketone biosynthesis in wheat (Simpson and von Wettstein-Knowles, 1980; Schneider *et al.*, 2016; Hen-Avivi *et al.*, 2016). However, upstream regulation of the expression and activity of these components of the wax pathway remains little understood (von Wettstein-Knowles, 2017).

In this paper, we describe how we mobilized advanced genotyping technologies and exploited germplasm resources at NordGen and USDA-ARS to identify the gene underlying the *Cer-X* locus controlling the wax bloom. We show that *Cer-X* encodes a SHINE family transcription factor, HvWAX INDUCER1 (HvWIN1), that we propose has a regulatory role in promoting cuticle integrity and wax bloom formation. We also show that while HvWIN1 is largely conserved among barley genotypes, two non-synonymous variants are specific to cultivated barleys.

## Materials and Methods

### Plant material and growth conditions

All barley *(Hordeum vulgare)* parent cultivars (*cv*.) and mutants are listed in Table S1. The *cer-x* alleles and their backgrounds, *cv*.’s Bonus, Nordal, Carlsberg II, Foma and Kristina, were obtained from NordGen, Alnarp, Sweden (www.nordgen.org). *cv*. Gateway and *gsh4.l* were obtained from the National Plant Germplasm System (NPGS) at the U.S. Department of Agriculture-Agricultural Research Service (USDA-ARS) (www.ars.usda.gov). *cv*. Bowman and the Near Isogenic Lines were obtained from the James Hutton Institute (Druka *et al.*, 2011). Plants were grown in general purpose cereal compost under long day glasshouse conditions (16 hours light 18°C/ 8 hours dark 14°C), with supplemental light as required.

### Cuticle integrity measurements

Cuticle integrity was assessed using chlorophyll leaching and Toluidine Blue (TB) staining assays. For chlorophyll leaching assays, fresh tissues were isolated from the second fully expanded leaf (growth stage 12, GS12; (Zadoks *et al.*, 1974)), flag leaf (GS55) and leaf sheath (GS55), with three (leaf sheath) or four (leaves) biological replicates per genotype. Fresh weight was recorded for each tissue sample before immersing samples in 80% ethanol and agitating them at 60rpm in the dark at room temperature. Following tissue immersion, absorbance of the ethanol solution was measured with a Varioskan Lux using Skanit microplate reader software (ThermoFisher technologies) every hour for 6 hours. A_664_ and A_647_ were recorded as maximum absorbance of chlorophyll A and B respectively; A_720_ was used to subtract the background signal and A_900_ and A_975_ corrected for path length. Absorbance values were normalised to a 1cm pathlength using the formulae: path length = (A_977_-A_900_)/0.036, A_x_ (1cm) = (A_X_-A_720_)/path length (where A_X_ is the absorbance to be corrected) (Warren, 2008; Lampinen *et al.*, 2012). Total chlorophyll concentration was then calculated using the formula: total chlorophyll (μg/mL)= 7.93 x (A_664_) + 19.53 (A_647_) (Lolle *et al.*, 1997). A sample of 80% ethanol was used as blank. Data were normalised to fresh weight. Area under the curve was calculated for each genotype and unpaired student *t*-tests were used to check for statistical significance between genotypes. Data were plotted using the ggplot2 R function (Wickham, 2016). TB staining was used to examine cuticle permeability in leaf, leaf sheath and spike. Samples were immersed in 0.05% (w/v) aqueous TB for 5 hours (spike) or 24 hours (leaf and leaf sheath), rinsed with water and photographed.

### Genotyping and candidate gene sequencing

BW407 and BW126, along with Bowman, Bonus and Gateway, were genotyped using the barley 50K iSelect SNP chip (Bayer *et al.*, 2017). Markers were ordered based on barley *cv.* Morex genome assembly (Morex V3 (Mascher *et al.*, 2021)) and plotted to visualise their positions along the barley chromosomes using the ggplot2 R function (Wickham, 2016). *HvWIN1* (HORVU6Hr1G038120) was amplified using GoTaq DNA Polymerase with 5X Colorless GoTaq Reaction Buffer (Promega). Primer sets and PCR cycle conditions are listed in Table S2. The amplicons were cleaned with ExoSAP (Applied Biosystems) and sequenced. The *HvWIN1* gene model was plotted using the ggplot2 R function (Wickham, 2016).

### Gene expression

Hull tissues at 5 days post anthesis (DPA) were harvested, snap-frozen and ground to a fine powder in liquid nitrogen. Total RNA was isolated using Qiagen RNeasy Plant Mini Kit and cDNA was synthesised using ProtoScript II First Strand cDNA Synthesis Kit (New England Biolabs). SYBR Green Power Up (Thermofisher) was used to measure transcript levels of *HvCER-C, HvCER-Q, HvCER-U, HvCER1, HvCER1.2, HvWAX ESTER SYNTHASE/DIACYLGLYCEROL ACYLTRANSFERASE 1 (HvWSD1), HvLONG-CHAIN ACYL-COA SYNTHETASE 2 (HvLACS2), HvKCS1* and *HvKCS6.* Primers for *HvCER-C, HvCER-Q* and *HvCER-U* were taken from (Hen-Avivi *et al.*, 2016), primers for *HvCER1, HvKCS6* and *HvWSD1* were taken from (Duan *et al.*, 2015), primers for *HvKCS1* were taken from (Li *et al.*, 2018), and primers for *HvLACS2* were taken from (Kumar *et al.*, 2016). qRT-PCR was as in (Shoesmith *et al.*, 2021) with *ACTIN7* (*HvACT7*) as an endogenous control. Primer sets are listed in Table S2. Data were plotted using the ggplot2 R function (Wickham, 2016). We searched the Barley Expression Database (EoRNA, (Milne *et al.*, 2021)) using the corresponding barley gene reference transcript model (BART1_0-p44305; BaRTV1.0, (Rapazote-Flores *et al.*, 2019)) to profile *HvWIN1* expression across 16 different tissues in Morex.

### Wax quantification

Flag leaves, flag leaf sheaths and spikes at GS55 were harvested and immediately frozen at −80°C. Surface waxes were extracted from one spike, one flag leaf and one 10 cm segment of leaf sheath by dipping the sample for 1 minute in 20mL dichloromethane containing 10μg methyl-nonadecanoate as the internal standard and dried under a vacuum evaporator. Extracts were derivatised by resuspending in 200μl (spikes and leaf sheaths) or 100 μl (leaves) N-O-bis-trimethylsilyltrifluoroacetamide (BSTFA) and incubating at 140°C for 1 hour. Wax components were identified using Gas Chromatography-Mass Spectrometry (GC-MS) as in (Brennan *et al.*, 2017) with the following modifications: the programmable temperature vaporising (PTV) injector operated in split mode (40:1 ratio), and solvent delay for mass spectrum acquisition was 2.8 min. Data were acquired and analysed using Xcalibur™ (version 2.0.7, Thermo Fisher Scientific): specific ions, characteristic of each compound, were selected and used for both compound detection and quantification in a processing method. Processed data were manually checked and corrected where necessary. Compounds that were found in less than three (out of four) replicates were assigned as “traces”. Data were plotted using the ggplot2 R function (Wickham, 2016).

### Sequence retrieval and phylogeny

We identified SHINE proteins by conducting a BLASTP search with the HvWIN1 protein sequence against the following databases. *Physcomitrium patens, Selaginella moellendorffii, Marchantia polymorpha, Amborella trichopoda, Arabidopsis thaliana, Solanum lycopersicum, Solanum tuberosum, Brassica rapa, Oryza sativa Japonica, Zea mays, Sorghum bicolor, Brachypodium distachyon, Triticum aestivum* and *Hordeum vulgare* sequences were obtained from EnsemblPlants (https://plants.ensembl.org/ (Bolser *et al.*, 2016)), *Pinus sylvestris* and *Picea abies* from PLAZA Gymnosperms (https://bioinformatics.psb.ugent.be/plaza/versions/gymno-plaza/), and *Azolla filiculoides* and *Salvinia cucullata* from FernBase (https://www.fernbase.org/). Additionally, the SlSHN2 (Solyc12g009490.1.1) protein sequence was obtained from Sol Genomics (https://solgenomics.net/ (Fernandez-Pozo *et al.*, 2015)) and Traes_6DS_E6A0BE6CD protein sequence was obtained from Phytozome (Goodstein *et al.*, 2012) (https://phytozome-next.jgi.doe.gov/). Sequences with apparent assembly errors were manually corrected by genomic sequence comparison against other putative SHINEs. Orthologues were selected based on the presence of three conserved motifs (AP2 domain, middle motif and c-terminal motif) and further explored using MEME motif discovery (http://meme-suite.org/tools/meme). Site distribution was set to one occurrence per sequence (OOPS) for a maximum of six motifs per sequence with motif widths of 5-70 amino acids (Bailey and Elkan, 1994).

Full protein sequences were aligned in MEGA 11 using MUSCLE. The evolutionary history was inferred by using the Maximum Likelihood method and JTT matrix-based model (Jones *et al.*, 1992) and tested using 300 bootstrap replications. Initial tree(s) for the heuristic search were obtained automatically by applying Neighbor-Join and BioNJ algorithms to a matrix of pairwise distances estimated using the JTT model, and then selecting the topology with superior log likelihood value. A discrete Gamma distribution was used to model evolutionary rate differences among sites (5 categories (+G, parameter = 0.7322)). The tree is drawn to scale, with branch lengths measured in the number of substitutions per site. This analysis involved 43 amino acid sequences. All positions containing gaps and missing data were eliminated (complete deletion option) resulting in 149 positions in the final dataset. Evolutionary analyses were conducted in MEGA11 (Tamura *et al.*, 2021).

### Haplotype analysis

SNP data of *HvWIN1* was retrieved from published exome-capture datasets of *H. spontaneum* and *H. vulgare* lines (Russell *et al.*, 2016; Bustos-Korts *et al.*, 2019). The dataset collected included 3 Kb upstream and 3 Kb downstream of the coding region and was filtered to retain sites with ? 98% of samples homozygous. Accessions with missing data points or heterozygosity at these sites were excluded. The resulting dataset, including 456 accessions, was used to build two *HvWIN1* haplotype networks, one containing only exonic SNPs and another containing all SNP sites in the dataset, including the 3 Kb upstream and downstream. Median-Joining haplotype network construction was performed using *PopArt (popart.otago.ac.nz* (Leigh and Bryant, 2015)). Exonic site haplotypes were plotted on a world map using the rworldmap R package (v1.3-6; (South, 2011)). Where available, latitude and longitude of sampling or based on sub-national production mid-point as in (Russell *et al.*, 2016; Bustos-Korts *et al.*, 2019) were used. If unknown, the latitude and longitude of the capital city of the country of origin were used. This information was not available for nine lines which were therefore excluded from this analysis. In addition, *HvWIN1* sequences from 19 barley accessions were retrieved from published sequencing data (Jayakodi *et al.*, 2020). Full length genomic sequences, including 3 Kb upstream and 3 Kb downstream, were aligned using ClustalW in MEGA-X (version 10.1.8, (Tamura *et al.*, 2021)), and the alignment used to identify SNPs and define haplotypes.

## Results

### Cer-X controls cuticular integrity and epicuticular wax composition

To investigate the mechanisms underlying cuticle formation in barley, we examined the *cer-x* Bowman Near Isogenic Line (Druka *et al.*, 2011), BW407, harboring the *glossy sheath4.l* (*gsh4.l*) allele originally identified in a radiation induced mutant in Gateway (Table S1). Compared to the glaucous appearance of Gateway, the *gsh4.l* mutant displays a striking glossy appearance on spikes, leaf sheaths and exposed nodes and internodes, characteristic of a loss of crystalline epicuticular waxes (Lundqvist and Wettstein, 1962; Rasmusson and Lambert, 1965; Mikkelsen, 1979). We confirmed that BW407 exhibits glossy phenotypes similar to the original *gsh4.l* allele and in contrast with the glaucous recurrent parent Bowman (Figure 1A-C, Table S3).

**Figure 1.**
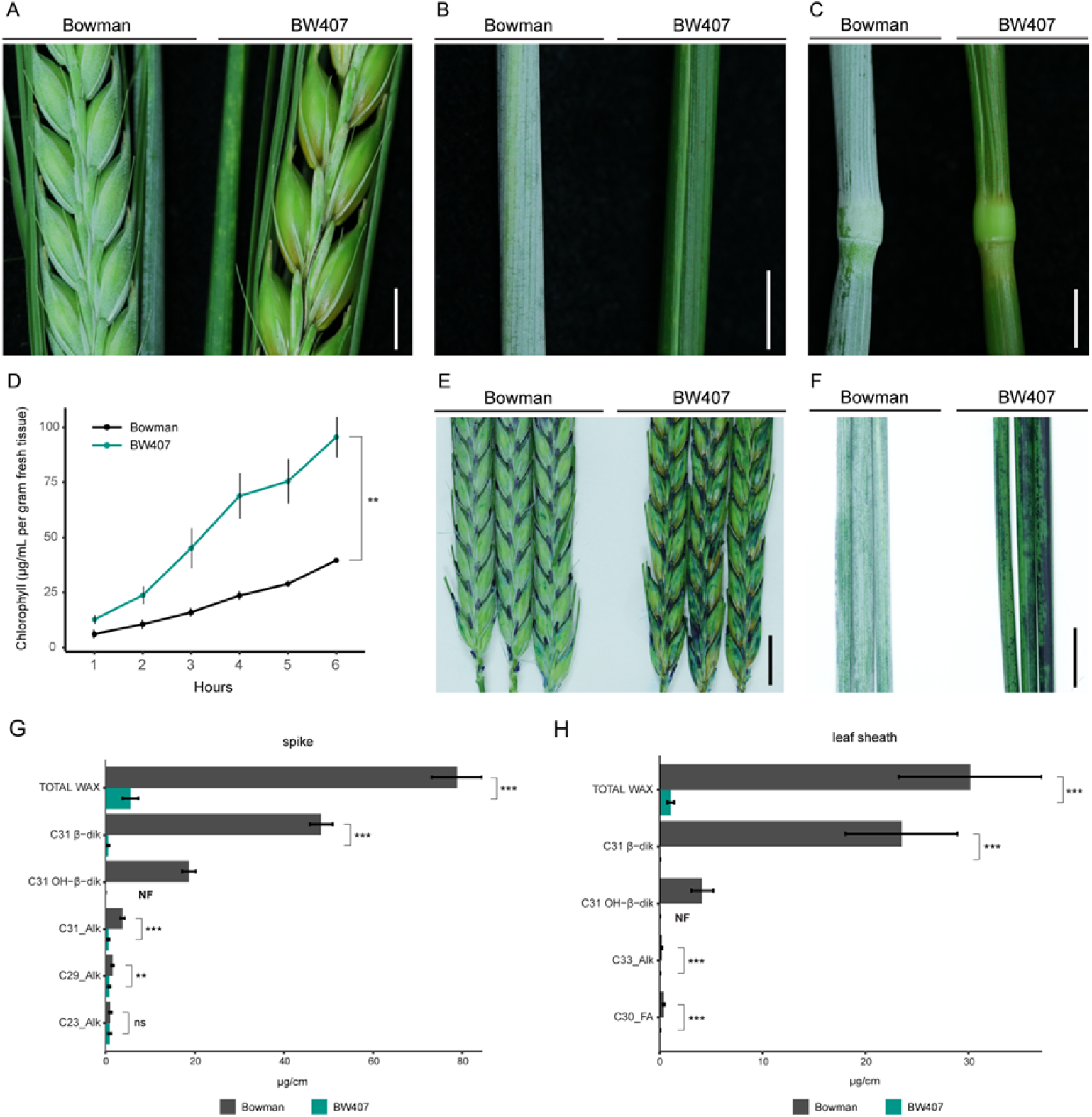
The *Cer-X* locus regulates specific cuticle structural and chemical properties in barley. (A-C) Wax crystal deposition on spikes, flag leaf sheaths and stems (nodes and internodes) of Bowman and BW407. Scale bars, 5 mm. (D) Chlorophyll leaching of Bowman and BW407 flag leaf sheaths. Bars represent standard error (n=3). *P*=0.00145 (unpaired student *t*-test). (E-F) Toluidine blue staining of spikes and flag leaf sheaths of Bowman and BW407. Scale bars, 10 mm. (G-H) Quantification of surface wax components extracted from (G) spikes and (H) flag leaf sheaths at GS55. NF = not found. Alk=Alkane, FA=Fatty Acid; β-dik = β-diketones; OH-β-dik = OH-β-diketones. * *P* < 0.05, ** *P*< 0.01, *\*\*\* P* < 0.001 (twotailed student *t*-test).

We next examined whether the *gsh4.l* allele influences cuticular properties in tissues showing the glossy phenotype (flag leaf sheaths and spikes) as well as those without a visible phenotype (fully expanded leaf blades from the second leaf and flag leaf). We used chlorophyll leaching as a proxy for cuticle permeability, with an increased permeability associated with cuticular defects (Aharoni *et al.*, 2004). We first measured how readily chlorophyll leaches from BW407 versus Bowman tissues following immersion in 80% ethanol. Leaf blades showed no differences in chlorophyll leaching between genotypes but chlorophyll leached more quickly from leaf sheaths in BW407 compared to Bowman (*P*=0.00145; Figure 1D; Figure S1A,B), suggesting increased cuticular permeability in BW407. Next we tested cuticle integrity by immersing tissues in a toluidine blue (TB) solution which penetrates and stains tissues with discontinuous or defective cuticles (Tanaka *et al.*, 2004). Both spikes and leaf sheaths in BW407 compared to Bowman were more permeable to TB when immersed into a 0.05% TB solution for 5 hours and 24 hours, respectively, while no significant difference was observed in the leaf blades after 24 hours (Figure 1E,F; Figure S1C). Based on these data, we suggest that BW407 has weakened cuticle integrity on leaf sheaths and spikes compared to Bowman, but not in leaves. Thus, we conclude that the *Cer-X* locus is important for cuticle integrity in tissues which develop a wax bloom.

We next assessed whether decreases in the β-diketone aliphatics which make up the epicuticular wax tubes (Mikkelsen, 1979) could explain the loss of glaucous wax in BW407. We used gas chromatography-mass spectrometry (GC–MS) to perform partial compositional analysis on flag leaf, flag leaf sheath and spike wax extracts from BW407 and Bowman. As reported previously, leaf wax is predominantly composed of C_26_ alcohols, making up 69% and 71% of total leaf wax in Bowman and BW407, respectively. Neither total wax accumulation nor levels of identified single components differed between BW407 and Bowman leaves (Figure S1D; Table S4). In contrast, BW407 leaf sheaths and spikes showed 96% and 93% reductions in total wax, respectively, compared to Bowman. As expected, C_31_ β-diketones (β-diketones and OH-β-diketones) contribute 92% and 85% of the total wax from flag leaf sheaths and spikes, respectively, in Bowman, and while BW407 showed reductions in most wax components compared to Bowman, β-diketones show the greatest drop, accumulating to only 0.044% and 0.86% Bowman levels in BW407 flag leaf sheaths and spikes, respectively (Figure 1G,H; Table 1; Table S4; *P* < 0.001, *t*-test). We also detected lower levels of C_30_ FAs in leaf sheaths and reductions in several long chain alkanes in spikes (Table 1; Table S4). Higher residual β-diketones in BW407 spikes compared to leaf sheaths may reflect BW407’s glaucous spike rachis. Interestingly, while C_18_ FAs were not robustly detected in Bowman leaf sheaths, higher levels were detectable in BW407 leaf sheaths. Altogether, our evidence supports that loss of β-diketones leads to the loss of glaucousness in BW407 leaf sheaths and spikes, the same tissues showing compromised cuticular integrity.

**Table 1.** Relative abundance of major wax components on flag leaf sheaths and spikes ^a^

| Tissue | Compound <sup>b</sup> | Bowman <sup>c</sup><br>μg/cm<br>(% of total) | BW407 <sup>c</sup><br>μg/cm<br>(% of total) | Significance <sup>d</sup> |
| --- | --- | --- | --- | --- |
| Flag leaf sheath | $C_{33\_}Alk$ | 0.21 (0.71%) | 0.02 (1.53%) | *** |
| | $C_{31}$ β-diketone | 23.5 (77.91%) | 0.01 (1.14%) | *** |
| | $C_{30\_}FA$ | 0.39 (1.30%) | 0.02 (1.91%) | *** |
| | $C_{31}$ OH-β-diketone | 4.14 (13.71%) | NF | / |
|  | TOTAL WAX | 30.17 | 1.08 | *** |
| spike | $C_{23\_}Alk$ | 1.02 (1.29%) | 0.88 (17.1%) | ns |
| | $C_{29\_}Alk$ | 1.51 (1.91%) | 0.78 (15.2%) | ** |
| | $C_{31\_}Alk$ | 3.79 (4.81%) | 0.63 (12.22%) | *** |
| | $C_{31}$ β-diketone | 48.35 (61.34%) | 0.58 (11.26%) | *** |
| | $C_{31}$ OH-β-diketone | 18.67 (23.68%) | NF | / |
|  | TOTAL WAX | 78.82 | 5.16 | *** |

<sup>a</sup> Complete dataset in Table S4
<sup>b</sup> Alk=Alkane, FA=Fatty Acid
<sup>c</sup> means of four bioreplicates and percentage of the total wax. NF = not found.
<sup>d</sup> significance of a *t*-test (two-tailed distribution, two samples equal variance): ns = not significant, \*\*<0.01, \*\*\*<0.001

### Cer-X is HvWIN1

We exploited existing germplasm resources and applied high throughput genotyping platforms to identify the gene underlying the *Cer-X* locus. We used the barley 50k iSelect SNP chip (Bayer *et al.*, 2017) to genotype two *cer-x* BWNILs, the aforementioned BW407 (*gsh4.l*, generated in Gateway) (Rasmusson and Lambert, 1965) and BW126 which contains the *cer-x.60* allele originally generated in Bonus (Lundqvist and Wettstein, 1962). We identified Gateway alleles across chromosome 6H in BW407, consistent with previous mapping data (Shahla and Tsuchiya, 1990), but did not distinguish clear introgression borders. However, two Bonus introgressions across chromosome 6H in BW126 overlapped with the BW407 region containing Gateway alleles (Figure 2A; Table S5). The first introgression spans a region between 452-457 Mb and contains the *rob1* locus previously shown to be closely linked to the *gsh4.l* locus in BW407 (Lundqvist and Franckowiak, 1997). This introgression also includes the gene *HORVU6Hr1G038120 (HORVU.MOREX.r2.6HG0479430* from Morex V2 (Monat *et al.*, 2019)) encoding a homolog of the WAX INDUCER1/SHINE1 (WIN1/SHN1) APETALA2 (AP2) transcription factor controlling wax and cutin biosynthesis in Arabidopsis thaliana (Broun *et al.*, 2004; Aharoni *et al.*, 2004)(Kumar *et al.*, 2016) which was previously named *HvWIN1,* and associated with cuticular FA levels and resistance to fungal infection in barley (Kumar *et al.*, 2016). Given the glossy mutant phenotypes, we speculated that loss of function alleles in *HvWIN1* may underlie *cer-x* alleles.

**Figure 2.**
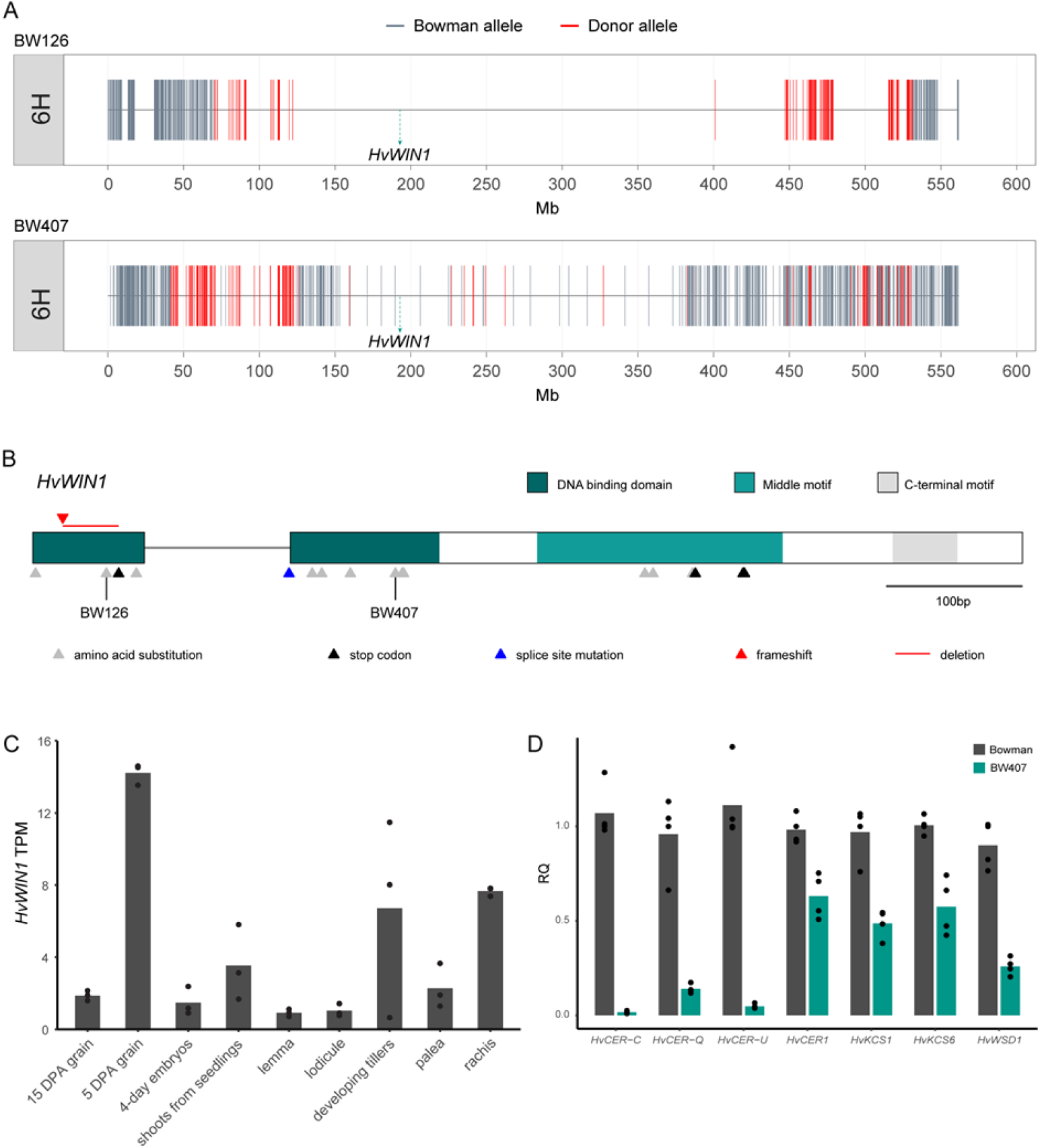
*HvWIN1* underlies the *Cer-X* locus in barley. (A) Barley 50k iSelect SNP chip (Bayer *et al.*, 2Ol7)plotted over chromosome 6H of BW126 and BW407. (B) *HvWIN1* gene model *(HORVU6Hr1G038120).* Variants in *cer-x* alleles represented as triangles. Triangles coloured based on mutation type. (C) Expression of *HvWIN1* in barley tissues from Barley Expression Database (EoRNA (Milne *et al.*, 2021)). Data expressed in transcripts per million (TPM). Bars represent mean expression. N=3. (D) Expression of HvWIN1 putative downstream targets in Bowman and BW407 hull tissues at 5 days post anthesis (DPA). Expressed as relative quantity (RQ). N=3.

Direct sequencing of *HvWIN1* in BW407 and BW126 identified unique point mutations in each line which cause single amino acid substitutions in the predicted AP2 domain of HvWIN1: M54R in BW407 and W19R in BW126 (Figure 2B; Table S3). These point mutations were confirmed in their donor mutants, *gsh4.l* and *cer-x.60,* compared to their parent cultivars (Table S3). Resequencing *HvWIN1* in 21 other independent *cer-x* alleles, which all exhibit loss or reduction in surface waxes, identified a further 17 unique mutations (Table S3). Of the 16 single point mutations identified, 11 caused single amino acid substitutions, four caused premature stop codons and one interrupted an intron splicing site (Figure 2B; Table S3). Another allele, *cer-x.407,* contained a 42 bp deletion causing a premature stop codon. All alleles impacted either the AP2 domain or the middle motif in the predicted protein, suggesting these regions are crucial to HvWIN1 function. *HvWIN1* transcripts are abundant in developing tissues including grain, tillers, and shoots from seedlings, as well as inflorescence tissues such as the rachis (Figure 2C; Figure S2A), supporting *HvWIN1’s* role in barley cuticle development. From these data, we conclude that variation in *HvWIN1* underlies the *cer-x* alleles and that *Cer-X* is *HvWIN1.*

### HvWIN1 influences gene expression associated with cuticle development

Since *HvWIN1* encodes a transcription factor, we speculated that loss of β-diketones and decreased cuticular integrity in *hvwin1* mutants reflected differences in *HvWIN1*-dependent gene expression. To explore this hypothesis, we compared the expression levels of selected candidate genes in Bowman and BW407, based on their demonstrated or predicted roles in wax synthesis and cuticle formation in barley. Consistent with β-diketone and OH-β-diketone deficiency, BW407 showed strikingly reduced *CER-C, CER-Q* and *CER-U* expression levels (Figure 2D). Expression of key genes controlling cuticular lipid biosynthesis were also reduced in BW407, including *HvKCS1* and *HvKCS6* (Figure 2D) which encode condensing enzymes producing VLCFAs, substrates which are necessary for various wax and cutin biosynthetic pathways, supporting our observation that most wax components are reduced in BW407 (Table S4). In addition, BW407 had decreased expression of *HvWAX ESTER SYNTHASE/DIACYLGLYCEROL ACYLTRANSFERASE 1 (HvWSD1),* a homolog of the Arabidopsis wax synthase *AtWSD1* involved in ester biosynthesis (Li *et al.*, 2008), and two homologs of alkane synthase *AtCER1* (Bourdenx *et al.*, 2011), *HvCER1* and *HvCER1.2* (Figure 2D; Figure S2B), consistent with the reduction of long chain wax esters and long chain alkanes in BW407 (Figure 1G; Table S4). We also detected slight reductions in the barley orthologue of *LONG CHAIN ACYL-COA SYNTHETASE 2 (LACS2;* Figure S2B) involved in generation of long chain acyl pools important for cutin biosynthesis and cuticle permeability in Arabidopsis (Schnurr *et al.*, 2004; Bessire *et al.*, 2007). Based on these results, we suggest that HvWIN1 may participate in a regulatory network to control the expression of genes important for cuticular integrity and the wax bloom in barley.

### HvWIN1 is part of a highly conserved gene family

The extent of glaucousness varies across grasses (Uddin and Marshall, 1988; Nice *et al.,* 2016; von Wettstein-Knowles, 2017; De la Fuente Cantó *et al.*, 2018; Würschum *et al.*, 2020; Qi *et al.*, 2021) with wild Triticeae species generally producing alcohol-rich wax at the reproductive stage (Tulloch *et al.*, 1980), suggesting that the β-diketone rich wax characteristic of cultivated wheat and barley may have been selected during domestication. To examine whether *HvWIN1* sequences show differences between the cultivated and wild germplasm, we compared whole-exome *HvWIN1* sequences from 456 wild (*H. spontaneum)* and cultivated (*H. vulgare)* barley accessions (Russell *et al.*, 2016; Bustos-Korts *et al.*, 2019). We identified only six variations in the coding region, all in the second exon of *HvWIN1,* suggesting that *HvWIN1* is highly conserved. Of these variants, two caused non-synonymous changes (D136N and T113I) in the SHINE-specific middle motif, with the T113I variant changing a highly conserved amino acid in SHINE orthologs across species. Variants arranged in seven haplotypes (HAP; Figure 3A; Table S6, Table S7). HAP1 and HAP2 contain over 90% of the germplasm (417 out of 456 accessions) and are the only two haplotypes represented across cultivars, landraces and wild barleys. HAP3, with 19 lines, was shared between landraces and wild barley, but excluded from cultivated barley. Interestingly, the two non-synonymous variants, T113I and D136N, were only found in HAP4 (13 sequences) and HAP5 (5 sequences), respectively, haplotypes made up of cultivars and landraces, excluding wild barley. Finally, HAP6 and HAP7 were found in one landrace and one wild barley, respectively. The geographical distribution of HvWIN1 haplotypes did not identify any strong association between a specific variant and its origin; however, all HAP5 lines and eight out of 13 HAP4 lines were collected from central, southern and eastern Asian countries, including Afghanistan, Pakistan, Tajikistan, Nepal, India and China, with a strong presence in regions of the Himalayan plateau (Figure S3B, Table S7). Analysis of the 3 Kb upstream and 3 Kb downstream of the *HvWIN1* coding sequence identified an additional 15 and 11 variants which formed 25 haplotypes, of which six included 90% of the lines (412 out of 456, Table S7; Table S8; Figure S3). We also compared *HvWIN1* genomic sequences across the barley pangenome, 19 barley accessions representative of global barley diversity (Jayakodi *et al.*, 2020). Similar to the previous dataset, we found very little variation in the coding region, with only two second exon SNP sites, both present in the previous dataset, including the T113I variant in two landraces: HvZDM01467 (also called Du-Li Huang or Dulihuang), one of the founders of the Chinese breeding program (Jayakodi *et al.*, 2020), and HvHOR7552, a landrace from Pakistan (Table S9; Table S7). Analysis of the 3 Kb upstream and 3 Kb downstream of the *HvWIN1* coding sequence in the pangenome lines identified an additional 15 and 10 SNPs respectively, and no big structural changes (such as large deletions or introgressions) in the regions putatively containing regulatory domains (Table S9). Collectively, these data suggest that HvWIN1 is broadly conserved in barley, although two minor haplotypes exclusive to landraces and cultivated barley show changes to conserved amino acids.

**Figure 3.**
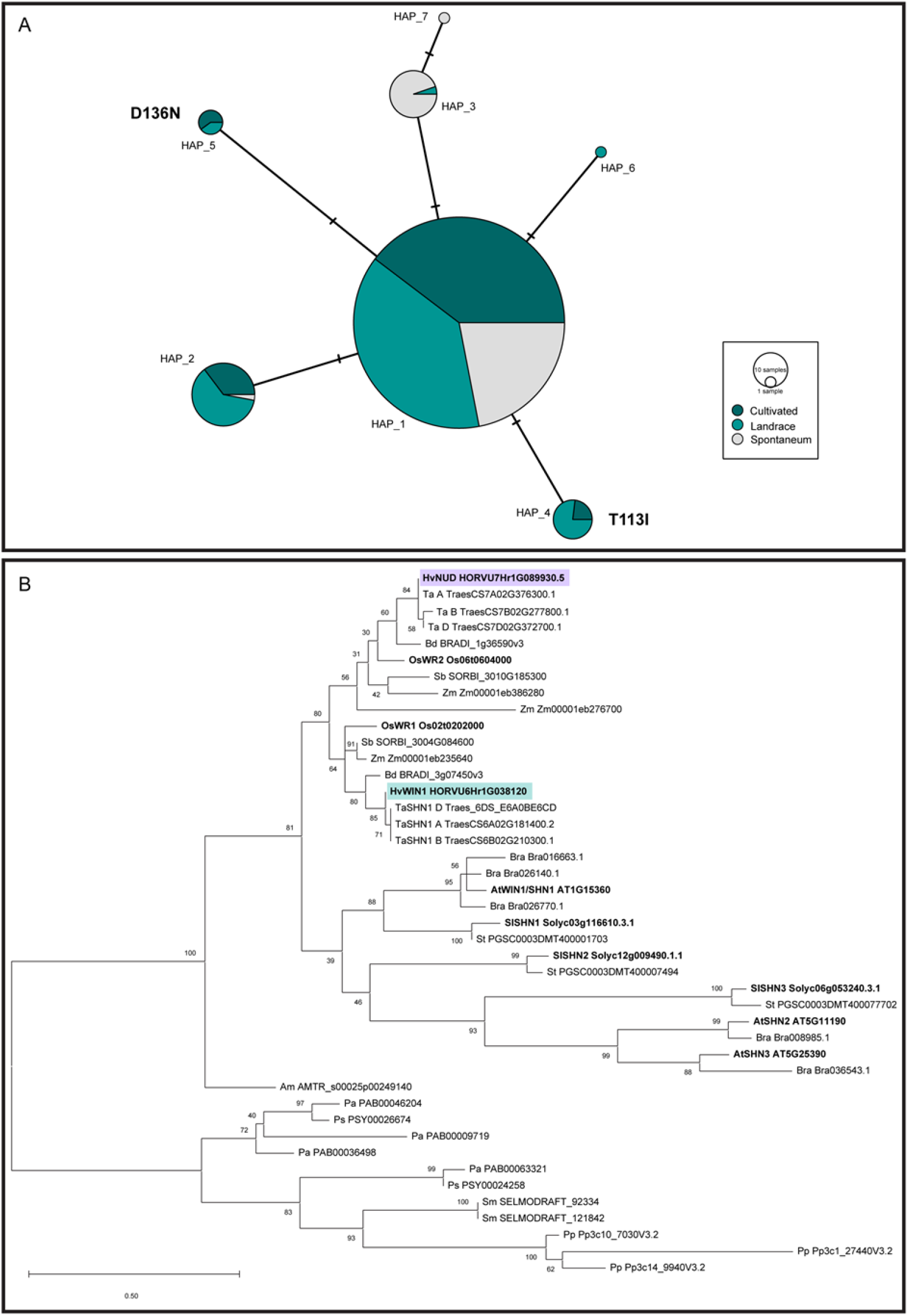
Sequence analyses of HvWIN1 and SHINE family transcription factors. (A) Median-joining network for *HvWIN1* haplotypes. SNPs were identified comparing *HvWIN1* exonic regions of 456 barley lines. Node size is relative to haplotype frequency. Bars between two nodes indicate the number of nucleotides within the sequence that differ between haplotypes. Amino acid changes are indicated in bold. (B) Phylogenetic relationship of SHINE family transcription factors across representative land plant species. Evolutionary analysis was inferred by using the Maximum Likelihood method and JTT matrix-based model in MEGA 11 (Jones *et al.*, 1992; Tamura *et al.*, 2021) The tree with the highest log likelihood (−3217.61) is shown. The percentage of trees in which the associated taxa clustered together in 300 bootstrap replications is shown next to the branches. Hv, *Hordeum vulgare;* Ta, *Triticum aestivum;* Bd, *Brachypodium distachyon;* Os, *Oryza sativa;* Sb, *Sorghum bicolor;* Zm, *Zea mays;* Bra, *Brassica rapa;* At, *Arabidopsis thaliana;* Sl, *Solanum lycopersicum;* St, *Solanum tuberosum;* Am, *Amborella trichopoda;* Pa, *Picea abies;* Ps, *Pinus sylvestris;* Sm, *Selaginella moellendorffii;* Pp, *Physcomitrium patens.* HvNUD is highlighted in purple, HvWIN1 is highlighted in green.

We next expanded our analyses to examine SHINE family transcription factors across representative species in the green plant lineage. Consistent with previous work (Kongeto/., 2020), SHINE proteins appear land plant-specific and likely emerged during plant adaptation to drier terrestrial environments, with SHINE homologs detected in the bryophyte *Physcomitrium patens* but not in other bryophytes or algal species (Figure 3B; Figure S4). SHINE homologs were highly conserved in all other land plants examined, with intriguing exceptions in water ferns. The closest homologs in the water ferns *Azolla filiculoides* and *Salvinia cucullata* contained partial SHINE motifs but low conservation of gene sequence and structure (Figure S4), though we note that lack of clear SHINE homologs in water ferns could also represent assembly errors. SHINE homologs form distinct subclades within both the dicot and monocot groups, suggesting that SHINE expansion and diversification events occurred separately in dicot and monocot lineages (Figure 3B). SHINE transcription factors act redundantly in floral organs in Arabidopsis (Shi *et al.*, 2011), but we observe neofunctionalisation in barley. The *NUDUM (HvNUD)* gene encodes another SHINE homolog whose deletion underlies ‘naked’ barley varieties where hulls shed freely due to a loss of a lipid-rich cementing layer on the grain pericarp which normally adheres to the hull (Taketa *et al.*, 2008). Comparing Bowman and the BWNIL introgressed with the *nud1.a* deletion (BW638) showed no change in leaf or leaf sheath cuticle permeability (Figure S5) and no visual wax phenotype (Table S3), suggesting that HvNUD does not regulate these traits, unlike HvWIN1.

## Discussion

The original *gsh4.l* and *cer-x* alleles were characterized by loss of surface wax coating on spikes, leaf sheaths and stems, which we confirmed reflect a loss of β-diketones, VLC alkanes and FAs in the BW407 *gsh4.l* introgression line. We also found that cuticular permeability increased in BW407. However, while total wax was reduced in BW407, changes in cuticular composition rather than total wax load may play more important roles explaining increased cuticular permeability in glossy BW407 tissues. Firstly, across multiple species and tissues, total wax content is not correlated to cuticle permeability (Jetter and Riederer, 2016; Seufert *et al.*, 2022). Secondly, completely β-diketone-deficient glossy *iw1Iw2* wheat lines also show increased chlorophyll leaching compared to glaucous β-diketone rich lines, despite equivalent total wax in both groups (Zhang *et al.*, 2013*b*). Taken together, our data is consistent with the role of β-diketones and alkanes in cuticular permeability in barley, which may underlie responses to drought and improved yield performance in glaucous varieties under arid conditions (Merah *et al.*, 2000; Zhang *et al.*, 2013*b*; Guo *et al.*, 2016; Bi *et al.*, 2017)

In addition to our study and to the best of our knowledge, the only other loss of function *shine* alleles reported are the *hvnud* alleles in barley (Taketa *et al.*, 2008) which showed a role for HvNUD in the control of cuticular lipids on the grain pericarp but not other cuticular surfaces. We show that HvWIN1 influences cuticular properties in both vegetative and reproductive tissues, consistent with previous work manipulating SHINE activity that showed roles in cuticular lipid metabolism as well as resistance to stress. In Arabidopsis, overexpression of *AtWIN1/SHN1* significantly increased wax and cutin accumulation and improved drought tolerance (Broun *et al.*, 2004; Aharoni *et al.*, 2004; Kannangara *et al.*, 2007) while silencing of a tomato *SHINE* gene *SlSHINE3 (SlSHN3)* caused reduced accumulation of fruit cuticular lipids, including cutin, and increased susceptibility to fungal infection and drought (Shi *et al.*, 2013). Similarly, transgenic knock-down of wheat *TaWIN1* reduced cuticular waxes and cutin, associated with increased susceptibility to fungal infection (Kong and Chang, 2018). Overexpression of rice *SHINE* orthologs, the wax synthesis regulatory gene 1 *(OsWR1)* and wax synthesis regulatory gene 2 *(OsWR2),* each increased drought tolerance and promoted wax synthesis; cutin levels also increased in the OsWR2 overexpression line, although these were not examined for *OsWR1* (Wang *et al.*, 2012; Zhou *et al.*, 2014). Barley spikes rub-inoculated with *HvWIN1* silencing constructs showed decreases in free FAs important for cutin biosynthesis and increased susceptibility to fungal infection, which the authors suggest reflects a role for HvWIN1 to reinforce the cuticle (Kumar *et al.*, 2016). Our data shows that variation in *HvWIN1* reduces cuticular waxes and alters cuticular integrity but whether HvWIN1 function also influences cutin levels in barley remains a pressing question. Moreover, we are curious whether non-synonymous cultivar-specific variation in *HvWIN1* have any influence on cuticular properties which could have been selected during cultivation, given the association between SHINE function and resistance to drought and infection.

Consistent with our work showing HvWIN1-responsive changes in cuticular metabolic gene expression, transgenic manipulation of SHINE function also suggested that SHINE transcription factors regulate expression of genes involved in both wax and cutin biosynthetic pathways (Broun *et al.*, 2004). Silencing of *SlSHN3* in tomato decreased expression of *SlCYP86A69,* a gene necessary for cutin accumulation in the tomato fruit cuticle (Shi *et al.*, 2013) while *OsWR1* overexpression and RNA interference lines in rice showed increased and decreased expression of *OsCER1, OsKCS1* and *OsCYP86A7* homologs, respectively (Wang *et al.*, 2012). *OsWR2* overexpression similarly promoted expression of *OsCER1, OsKCS1, OsLACS1* and *OsCYP86A7* homologs (Zhou *et al.*, 2014). Silencing *HvWIN1* in barley was also correlated with reduced expression of *HvCYP86A2, HvCYP89A2* and *HvLACS2* genes (Kumar *et al.*, 2016). Overexpressing *AtWIN1/SHN1* rapidly and directly induces the expression of cutin biosynthetic genes such as *AtCYP86A7, AtCYP86A4* and *AtLACS2* (Kannangara *et al.*, 2007), while wax biosynthetic genes such as *AtKCS1* and *AtCER1* are induced more slowly, suggesting that they operate further downstream (Kannangara *et al.*, 2007). We speculate that HvWIN1 may also control different gene targets depending on developmental stage and tissue. In barley, cutin and wax deposition reduced cuticular permeability in expanding leaves but further decreases in cuticular permeability reflected additional wax deposition in cells at their final length (Richardson *et al.*, 2007), suggesting carefully coordinated metabolic gene expression linked to differentiation. Moreover, environmental signals may be involved as deposition of epicuticular waxes appears linked to exposure to the atmosphere rather than age per se (Richardson *et al.*, 2005). Furthermore, barley and wheat leaf wax changes from predominantly primary alcohols and alkanes in seedlings to β-diketones in reproductive stage tissues such as sheaths, stems and spikes (von Wettstein-Knowles, 1972; Tulloch, 1973; Kosma and Jenks, 2007; Zhang *et al.*, 2013*b*). Learning more about HvWIN1 control of metabolic gene expression during development will be important to determine how HvWIN1 may control individual events and/or responses to environmental signals during vegetative and reproductive development to regulate cuticular integrity and cuticular waxes in specific tissues.

In summary, we discovered that variation in *HvWIN1* underlies alleles at the *Cer-X* locus which cause changes in cuticular integrity and cuticular waxes in barley. Our work was accelerated by a synergistic combination of access to invaluable germplasm resources and development of advanced genotyping platforms. Unlike most other genes so far identified which control cuticular waxes in barley, HvWIN1 encodes a transcription factor. Learning more about the regulatory network controlling cuticular features and how this may differ between cultivars and wild species may become increasingly important to develop more resilient cereal varieties better equipped to respond to environmental challenges.

## Supplementary Materials

Figure S1. The *Cer-X* locus does not affect leaf cuticle and wax properties in barley.

Figure S2. Expression of *HvWIN1* in barley tissues from Barley Expression Database (EoRNA).

Figure S3. Sequence variation of HvWIN1 in barley.

Figure S4. Motif analysis of SHINE family transcription factors.

Figure S5. The *NUD* locus does not affect leaf and leaf sheath cuticle properties in barley.

Supplemental Tables.xlsx

Table S1. Barley germplasm.

Table S2. Primer sequences.

Table S3. *cer-x* allele resequencing and visual scoring of wax coverage.

Table S4. Relative abundance of wax components on flag leaves, flag leaf sheaths and spikes.

Table S5. Genotyping data of Bowman, BW407, BW126, Gateway and Bonus using barley 50k iSelect SNP chip.

Table S6. Exonic haplotypes of *HvWIN1* discovered from 456 wild (*H. spontaneum)* and cultivated (*H. vulgare*) barley accessions.

Table S7. Lines used for *HvWIN1* haplotype analysis

Table S8. Up and downstream and whole genomic haplotypes of *HvWIN1* discovered from 456 wild (*H. spontaneum*) and cultivated (*H. vulgare*) barley accessions.

Table S9. Haplotypes of *HvWIN1* in the barley pangenome

## Acknowledgments

We are deeply grateful to the NordGen, National Plant Germplasm System at the U.S. Department of Agriculture-Agricultural Research Service and the James Hutton Institute for germplasm. We also acknowledge guidance from Dr. Alexandre Foito in generation of GC-MS data and Dr. Joanne Russell in haplotype analysis.

## Author Contributions

Conceptualization, S.M.M., C.C., and T.M.; methodology, S.M.M., C.C., and T.M; formal analysis, T.M. and C.C. ; investigation, T.M., C.C., L.L. and M.E.; writing—original draft preparation, S.M.M. and T.M.; writing—review and editing, S.M.M., C.C., and T.M.; supervision, S.M.M and C.C..; project administration, S.M.M..; funding acquisition, S.M.M. and C.C.. All authors have read and agreed to the published version of the manuscript.

## Conflicts of Interest

The authors declare no conflict of interest.

## Funding

This research was funded by the Biological and Biotechnological Research Council grant number BB/R010315/1 to S.M.M. C.C. and M.E. were supported by BB/R010315/1. M.E. was also supported by the University of Dundee and the Council for Academics at Risk (Cara). L.L. was supported by the China Scholarship Council and the University of Dundee. T.M. was supported by a Carnegie-Cant-Morgan PhD Scholarship and the University of Dundee.

## Data Availability Statement

Large datasets were not generated; however, R scripts were developed and will be deposited upon manuscript publication.

## Notes

### Competing Interest Statement

The authors have declared no competing interest.

