## Supplementary figures for "A WAX INDUCER1/SHINE transcription factor controls cuticular wax in barley"

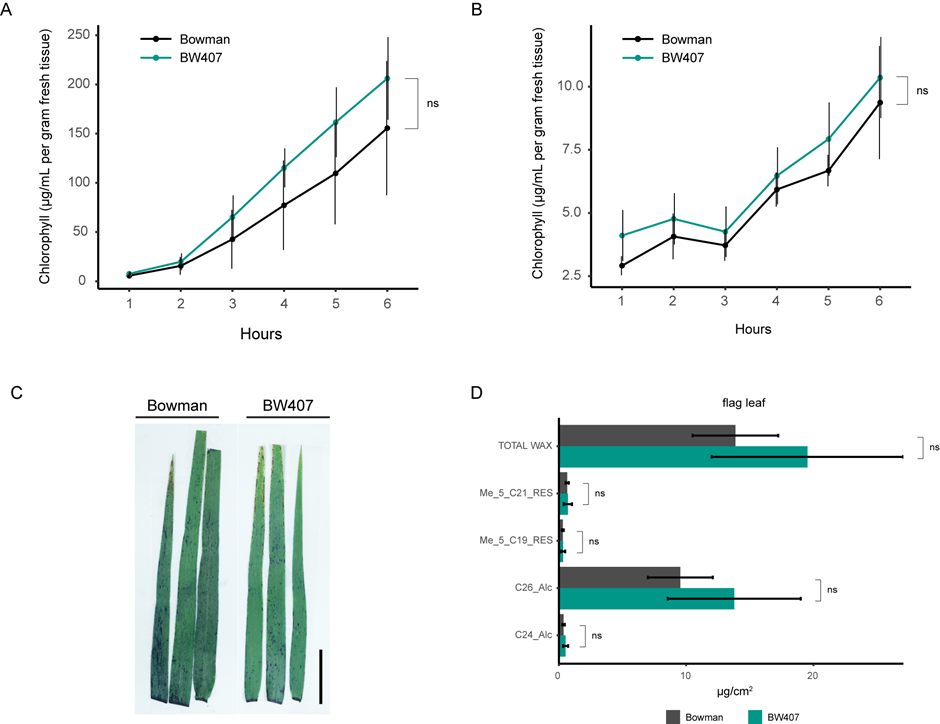

Figure S1. The *Cer-X* locus does not affect cuticular and wax properties in barley leaf blades. (A-B) Chlorophyll leaching of Bowman and BW407 (A) flag leaf (GS55) and (B) second leaf (GS12). Bars represent standard error. N=4. ns = not significant (unpaired student *t*-test). (C) Toluidine blue staining of flag leaves of Bowman and BW407. Scale bars, 20 mm. (D) Quantification of surface wax components extracted from flag leaves at GS55. Alc = alcohol, Me_5_RES = methyl alkyl resorcinol. ns = not significant (two-tailed student *t*-test). N=4.

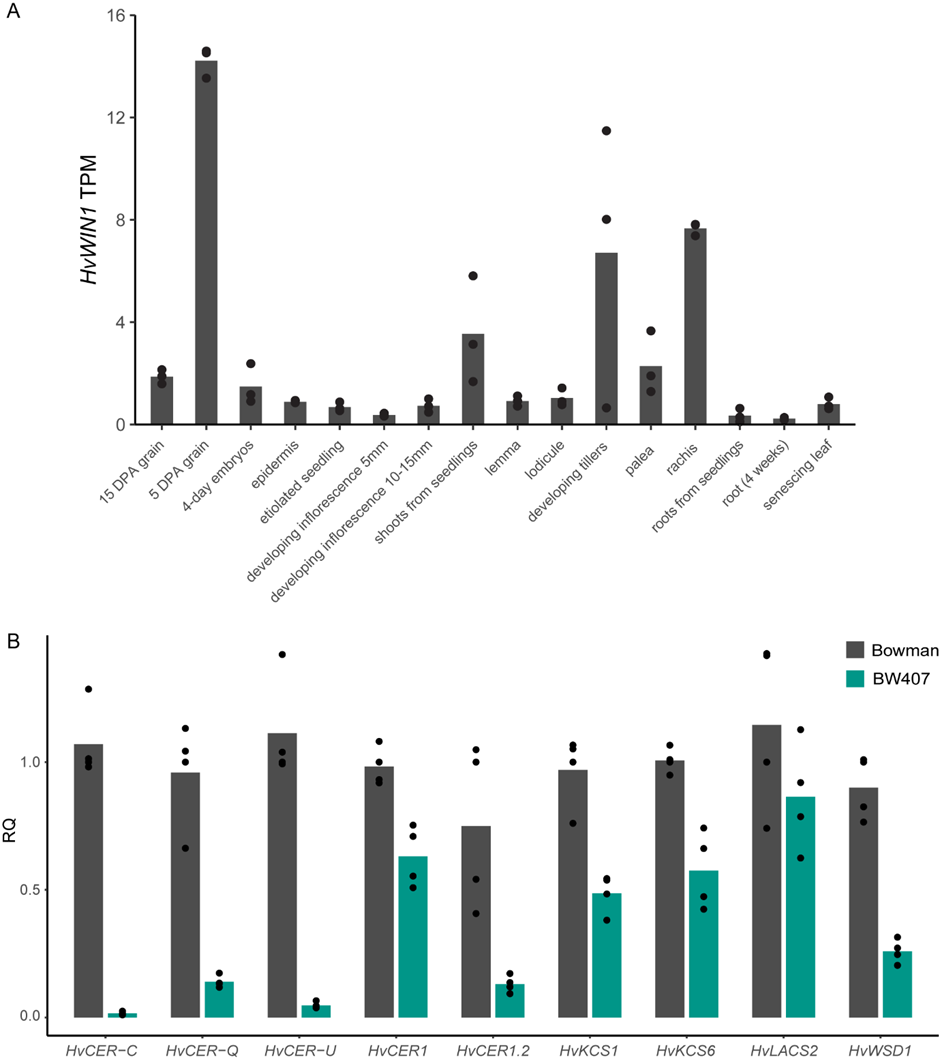

Figure S2. Full *HvWIN1* expression analyses. (A) Expression of *HvWIN1* in barley tissues from Barley Expression Database (EoRNA). Data expressed in transcripts per million (TPM). Bars represent mean expression. N=3. (B) Expression of *HvWIN1* downstream targets in Bowman and BW407 hull tissues at 5 days post anthesis (DPA). Expressed as relative quantity (RQ). Bars represent means of four bioreplicates (dots).

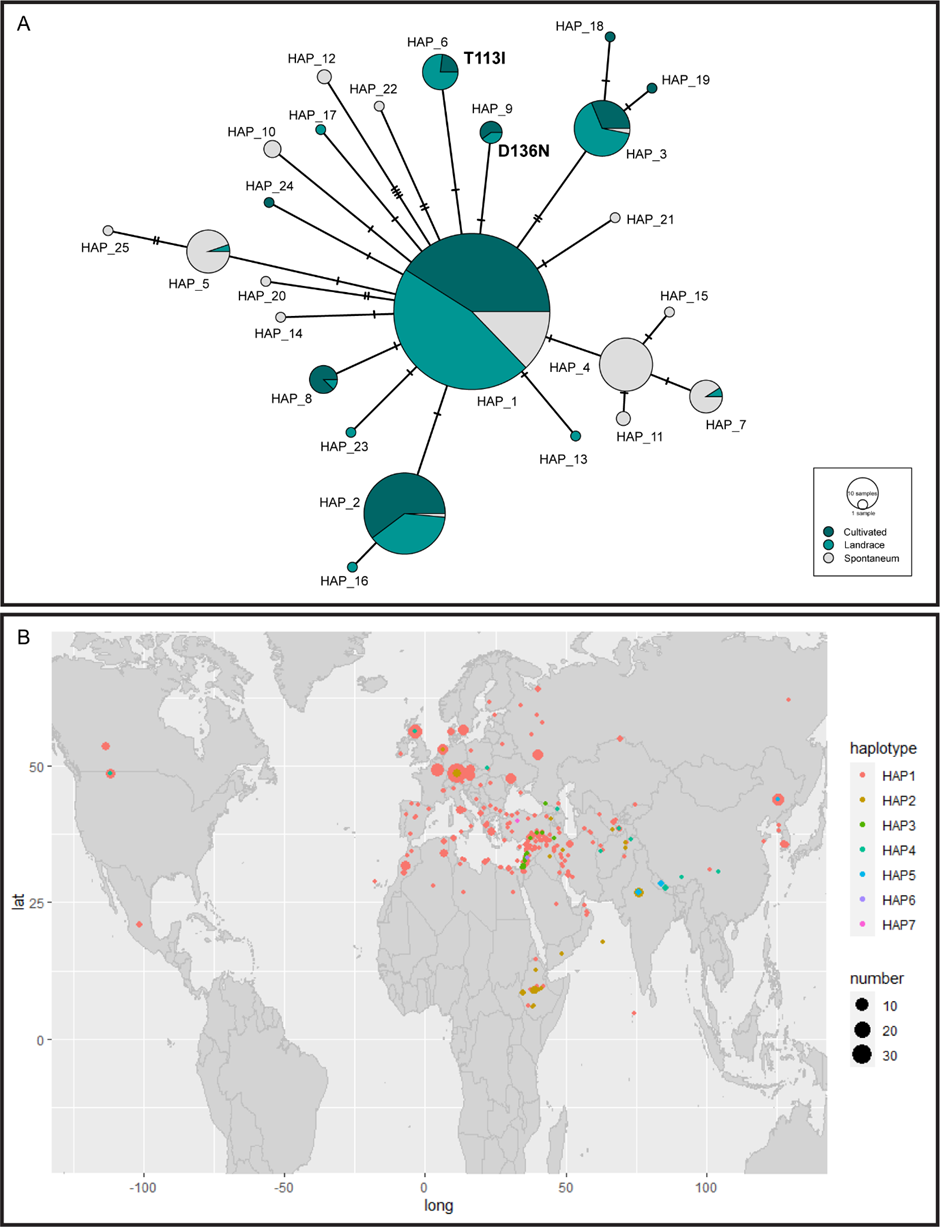

Figure S3. Sequence Variation of *HvWIN1* in barley. (A) Median-joining network for *HvWIN1* haplotypes. SNPs were identified comparing *HvWIN1* sequence including 3Kb up- and downstream the coding region, of 456 barley lines. Node size is relative to haplotype frequency. Bars between two nodes indicate the number of nucleotides within the sequence that differ between haplotypes. Amino acid changes are indicated in bold. (B) Geographical distribution of *HvWIN1* haplotypes defined by six SNP sites in the coding region. The size of the circle is proportional to the number of lines.

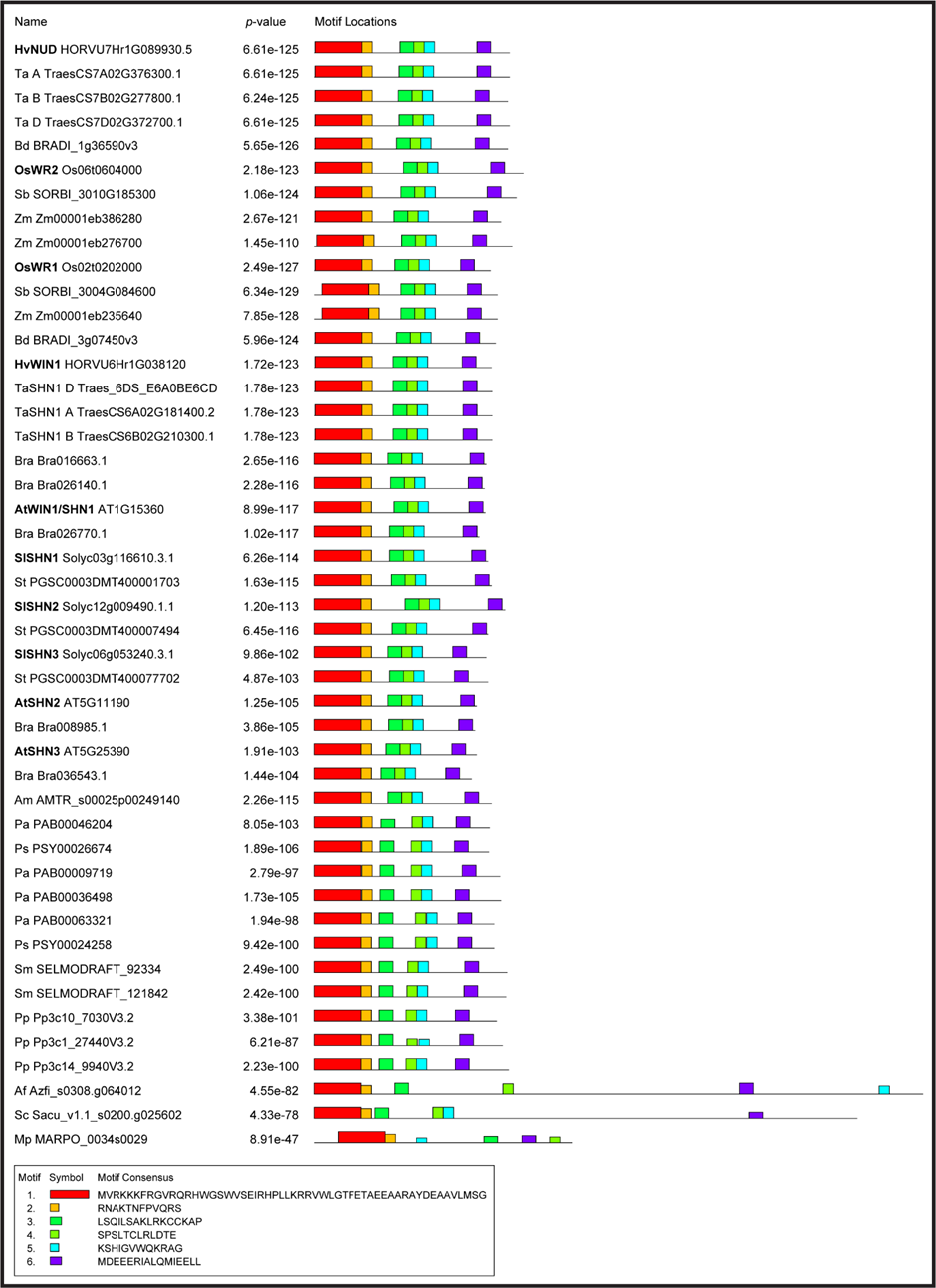

Figure S4. Motif analysis of SHINE family transcription factors. Top orthologues from a BLASTP using HvWIN1 were examined from representative species in the green plant lineage. SHINE orthologs were denoted by the presence of full SHINE motifs (AP2 domain [#1-2], middle motif [#3-5] and c-terminal motif [#6]). Consensus sequences for each motif are displayed at the bottom. Motif block height represents significance of the motif site, proportional to the negative logarithm of the *p*-value. Hv, *Hordeum vulgare*; Ta, *Triticum aestivum*; Bd, *Brachypodium distachyon*; Os, *Oryza sativa*; Sb, *Sorghum bicolor*; Zm, *Zea mays*; Bra, *Brassica rapa*; At, *Arabidopsis thaliana*; Sl, *Solanum lycopersicum*; St, *Solanum tuberosum*; Am, *Amborella trichopoda*; Pa, *Picea abies*; Ps, *Pinus sylvestris*; Sm, *Selaginella moellendorffii*; Pp, *Physcomitrium patens*; Af, *Azolla filiculoides*; Sc, *Salvinia cucullata*; Mp, *Marchantia polymorpha*.

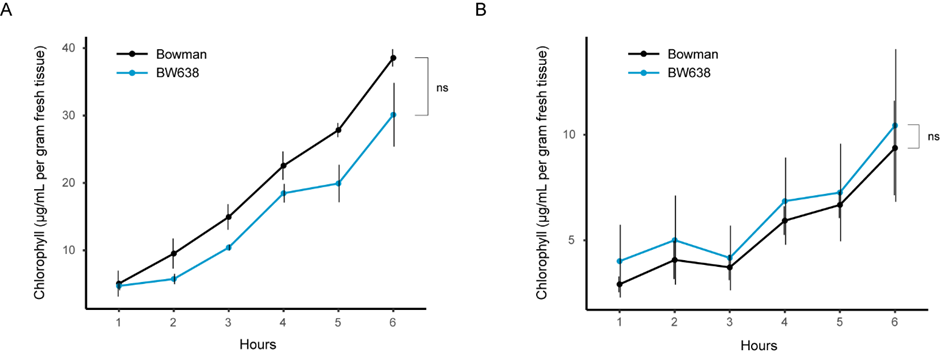

Figure S5. The *NUD* locus does not affect leaf and leaf sheath cuticle properties in barley. (A-B) Chlorophyll leaching of Bowman and BW638 (A) flag leaf sheath at GS55 and (B) second leaf at GS12. Bars represent standard error. N=3 in leaf sheath; N=4 in second leaf. ns = not significant (unpaired student *t*-test).
